# Cardiorenal and hepatic dysfunctions underlie metabolic alterations in the spinocerebellar ataxia type 7 mice

**DOI:** 10.64898/2026.01.14.699424

**Authors:** Marie-France Champy, Nadia Messaddeq, Ghina Bouabout, Chantal Weber, Anna Niewiadomska-Cimicka, Mohammed Selloum, Laurent Monassier, Bruno Moulin, Yvon Trottier

## Abstract

Spinocerebellar ataxia type 7 (SCA7) is a polyglutamine expansion disorder characterized by progressive cerebellar and retinal degeneration leading to ataxia and blindness. While mutant ATXN7 is ubiquitously expressed, most studies have focused on neurological symptoms, and peripheral contributions to pathology remain poorly understood. Here, we investigated systemic abnormalities in SCA7^140Q/5Q^ knock-in mice, a model of early-onset disease.

Using longitudinal metabolic profiling, ultrasound imaging, and histopathology, we identified early and progressive dysfunction of the kidney, liver, and heart. Renal impairment was marked by uremia, polyuria with abnormal calcium and glucose excretion, tubular epithelial cell loss, and podocyte dedifferentiation, including the reappearance of primary cilia and pedicel effacement. Hepatic alterations included dysregulated lipid metabolism, elevated bilirubin, and reduced iron levels, contributing to anemia. Cardiac dysfunction manifested as reduced stroke volume as early as 11 weeks, suggesting a cardiorenal syndrome. Together, these organ-specific changes resulted in systemic metabolic disturbances such as dyslipidemia, iron deficiency anemia, thrombocytosis, and chronic inflammation, detectable before the onset of motor incoordination.

Our findings demonstrate that peripheral organ dysfunction is an early and integral feature of SCA7 pathogenesis, with renal and cardiac impairments emerging prior to neurological decline. These results highlight the value of systemic biomarkers for disease monitoring and suggest that targeting peripheral pathology may provide therapeutic benefit. More broadly, they underscore the need to view SCA7 not solely as a neurodegenerative disorder but as a multi-organ disease.

## INTRODUCTION

Spinocerebellar ataxia type 7 (SCA7) is a rare autosomal dominant neurodegenerative disorder characterized by progressive cerebellar ataxia and retinal degeneration leading to blindness [1]. SCA7 is caused by abnormal expansion of CAG trinucleotide repeat in *ATXN7* gene, which encodes a polyglutamine (polyQ) tract in the protein [2]. Like Huntington’s disease (HD) and several other spinocerebellar ataxias (SCA 1-3, 6 and 17), SCA7 belongs to the group of CAG/polyQ expansion disorders [3]. Normal *ATXN7* alleles contain 4 to 35 CAG repeats, whereas pathological expansions usually exceed 36 and can reach several hundred [4,5]. Repeat length strongly influence disease onset and severity, with larger expansions typically causing severe early infantile and congenital forms leading to death within years or months [6]. These early-onset forms are characterized by hypotonia, failure to thrive and may develop extra-neural pathologies, affecting the kidney, liver and heart, which remain poorly characterized [5,6,7–14].

ATXN7 protein is widely expressed in both neural and non-neural tissues [15,16]. Within the nucleus, it is a key component of the Spt-Ada-Gcn5 Acetyltransferase (SAGA) complex, which regulates gene transcription through histone acetylation and deubiquitination [17,18]. Studies in zebrafish indicated that *Atxn7* plays a role in the terminal differentiation of photoreceptors and cerebellar Purkinje cells [19,20], two neuron types primarily affected in SCA7. PolyQ expansion in ATXN7 causes its aggregation and disrupts SAGA-mediated transcriptional control, leading to impaired differentiation and survival of photoreceptors and Purkinje cells [21–27]. SCA7^266Q/5Q^ and SCA7^140Q/5Q^ knock-in mouse models, which express mouse ATXN7 with early-onset allelic forms containing 266 and 140 glutamines, respectively, recapitulate disease features, exhibiting retinal and cerebellar degeneration and, in the latter, additional forebrain atrophy [25,28,29]. Notably, both models also show growth retardation, hypoactivity and premature death, the origins of which remain poorly understood. Increasing evidence from other polyQ disorders, such as Huntington’s disease, indicates that peripheral organs—including muscle, kidney, liver, and heart—can be directly affected [30,31]. However, the extent and significance of peripheral pathology in SCA7 remain largely unexplored.

In this study, we investigated peripheral organ involvement in the SCA7^140Q/5Q^ mouse model. Using ultrasound imaging, metabolic profiling and histopathology, we examined the kidney, liver and heart particularly affected in severe infantile/congenital SCA7 forms. Our findings reveal substantial pathological changes in these tissues that precede or accompany neurological decline, suggesting that peripheral dysfunction contributes to comorbidity and premature mortality in SCA7.

## RESULTS

### Plasma and urinary biochemistry of 18-week-old SCA7 mice

Earlier studies have shown that SCA7^140Q/5Q^ mice exhibit a gradual neurological phenotype, beginning with retinopathy from 10 weeks (wks) of age, motor incoordination at 20 wks and a median lifespan of 54 wks. Apart from having reduced body weight and shorter body length, SCA7^140Q/5Q^ mice did not exhibit other significant anatomical abnormalities [25]. At the end stage of disease (> 40wks), the proportions of lean, fat, and bone tissues were comparable to those of age-matched wild type (WT) mice, when normalizing to body weight [25]. There was no evidence of major damage to internal organs (spleen, heart, liver and kidney), and their weights normalized to body weight were comparable to those of WT littermates, except for kidneys that were 29% lighter than WT kidneys (p <0.0001) (Figure 1A-E).

**Figure 1:**
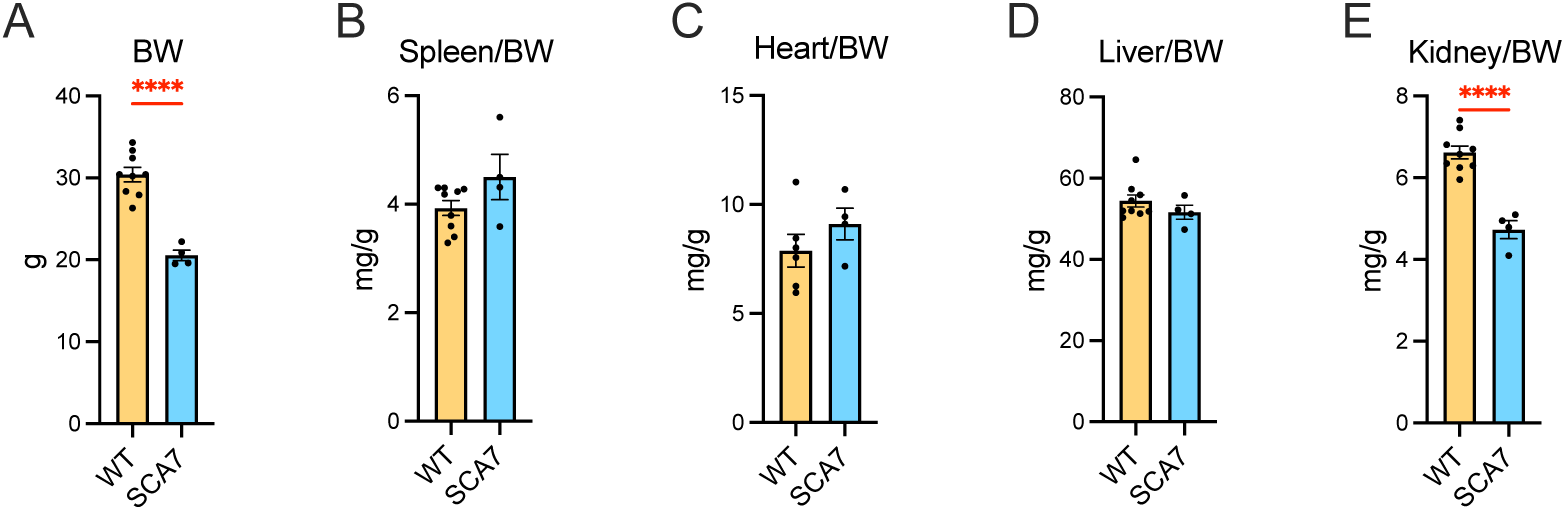
Post-mortem organ weights of SCA7 mice. The data show means ± SEM (n= 4 males/genotype). Body weight (BW). Means were analyzed using unpaired Student t-test. **** *p*< 0.0001.

### Plasma and urinary biochemistry of 18-week-old SCA7 mice

To evaluate metabolic and renal functions, we placed 18-wk-old SCA7 and WT male littermates in metabolic cages for a 24-hour biochemical analysis of plasma and urine. At this age, SCA7 males weighed 13% less than WT males (p< 0.0001) (Figure 2A). When normalized to body weight, SCA7 mice consumed 22% more food (p= 0.032) and 24% more water (p= 0.011), and excreted twice the urine volume (p= 0.0022) compared to WT mice (Figure 2B-D).

**Figure 2:**
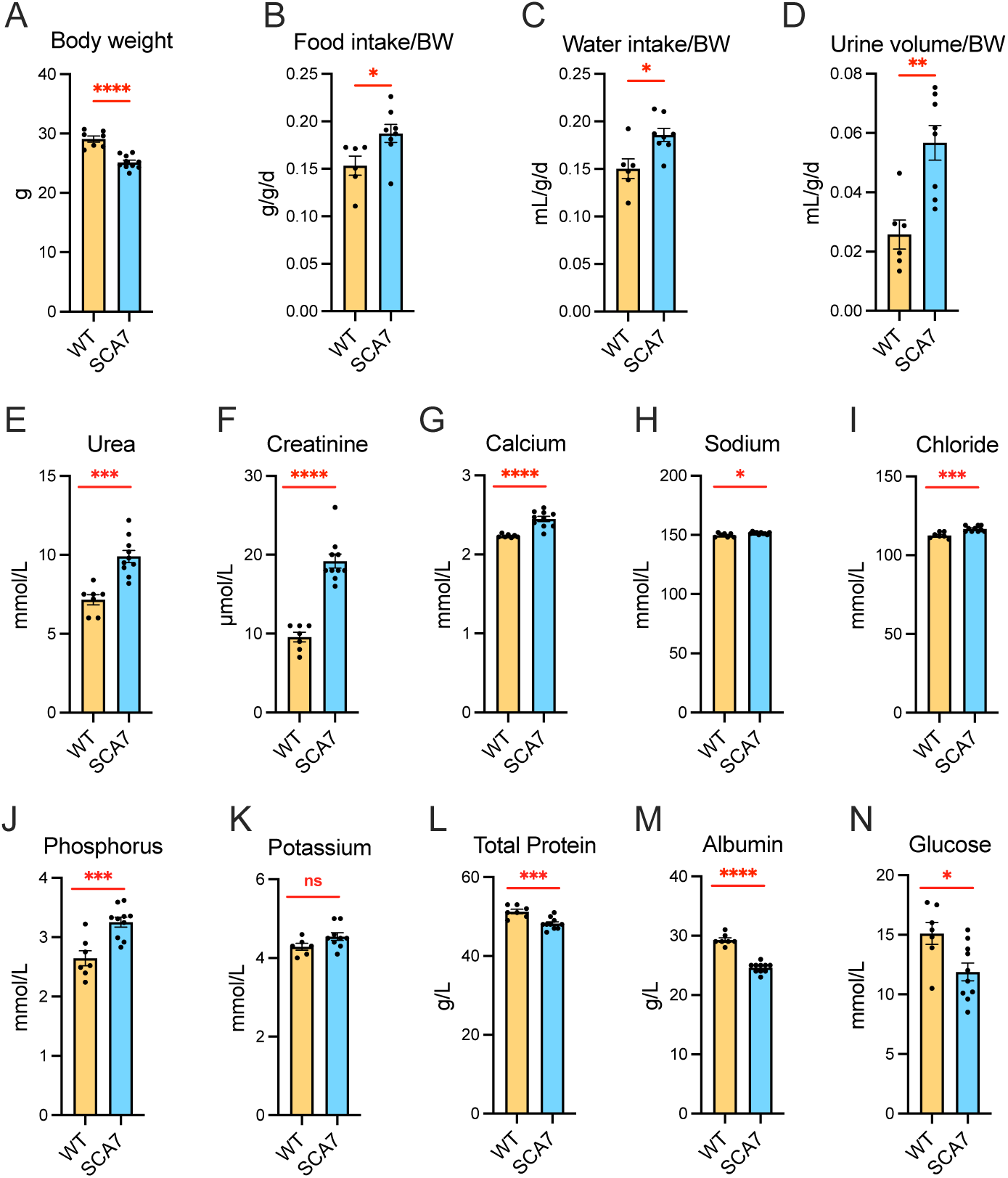
Plasma biochemistry parameters of SCA7 mice in metabolic cage. **A)** Body weight (BW) after 24 hours in metabolic cage of 18-wk-old SCA7 and WT male littermates. **B-D**) Total food intake, water intake, excreted urine volume during 24 hours normalized for BW. **E-N**) Concentrations of biochemical components in the plasma. Data are means ± SEM. *p* value compared WT (n= 7) versus SCA7 (n= 10) mice, using unpaired Student t-test. * *p* < 0.05, ** *p*< 0.01, *** *p*< 0.001, **** *p*< 0.0001. ns, not significant.

Plasma analysis revealed significantly elevated urea (+38%, p = 0.001) and creatinine (+100%, p < 0.0001) concentrations in 18-wk-old SCA7 mice (Figure 2E-F). Electrolyte levels were slightly but significantly increased, including calcium (+10%, p< 0.0001), sodium (+1%, p= 0.02), chloride (+4%, p= 0.0003) and phosphorus (+23%, p= 0.0008), whereas potassium remained unchanged (Figure 2G-K). In contrast, total proteins (-6%, p= 0.0007), albumin (-16%, p< 0.0001) and glucose (-21%, p= 0.015) concentrations were significantly reduced in SCA7 plasma (Figure 2L-N). Notably, the reduced glucose levels indicate that polydipsia and polyuria are not a consequence of diabetes mellitus.

These elevated metabolites and electrolytes, along with reduced total proteins, albumin and glucose levels, persisted in SCA7 plasma at 30 wks and at the end-disease stage of 43 wks, compared to age-matched WT (Supplementary Figure S1A-K). Analysis at an earlier age (16 wks) showed that plasma concentration of creatinine, total protein and albumin were the first parameters significantly affected in SCA7 males as well as in SCA7 females (Supplementary Figure S1L-N).

Urine analysis showed marked reductions in creatinine clearance (-50%, p< 0.0001) and sodium filtered load (-49%, p< 0.0001), indicating substantial renal impairment in 18-wk-old SCA7 mice (Figure 3A-B). SCA7 24-hour urine contained elevated levels of urea (+42%, p= 0.011), glucose (+306%, p= 0.020), calcium (+277%, p< 0.0001), sodium (+27%, p= 0.046) and phosphorus (75%, p= 0.026) (Figure 3C-I). However, consistent with the increased urine volume, osmolality was reduced by 39% (p< 0.0016), reflecting more dilute urine compared to WT controls (Figure 3J). Notably, urinary total protein (-68%, p= 0.0009) and albumin (-45%, p= 0.06) were decreased, suggesting that kidney dysfunction at this age is not associated with proteinuria (Figure 3K-L).

**Figure 3:**
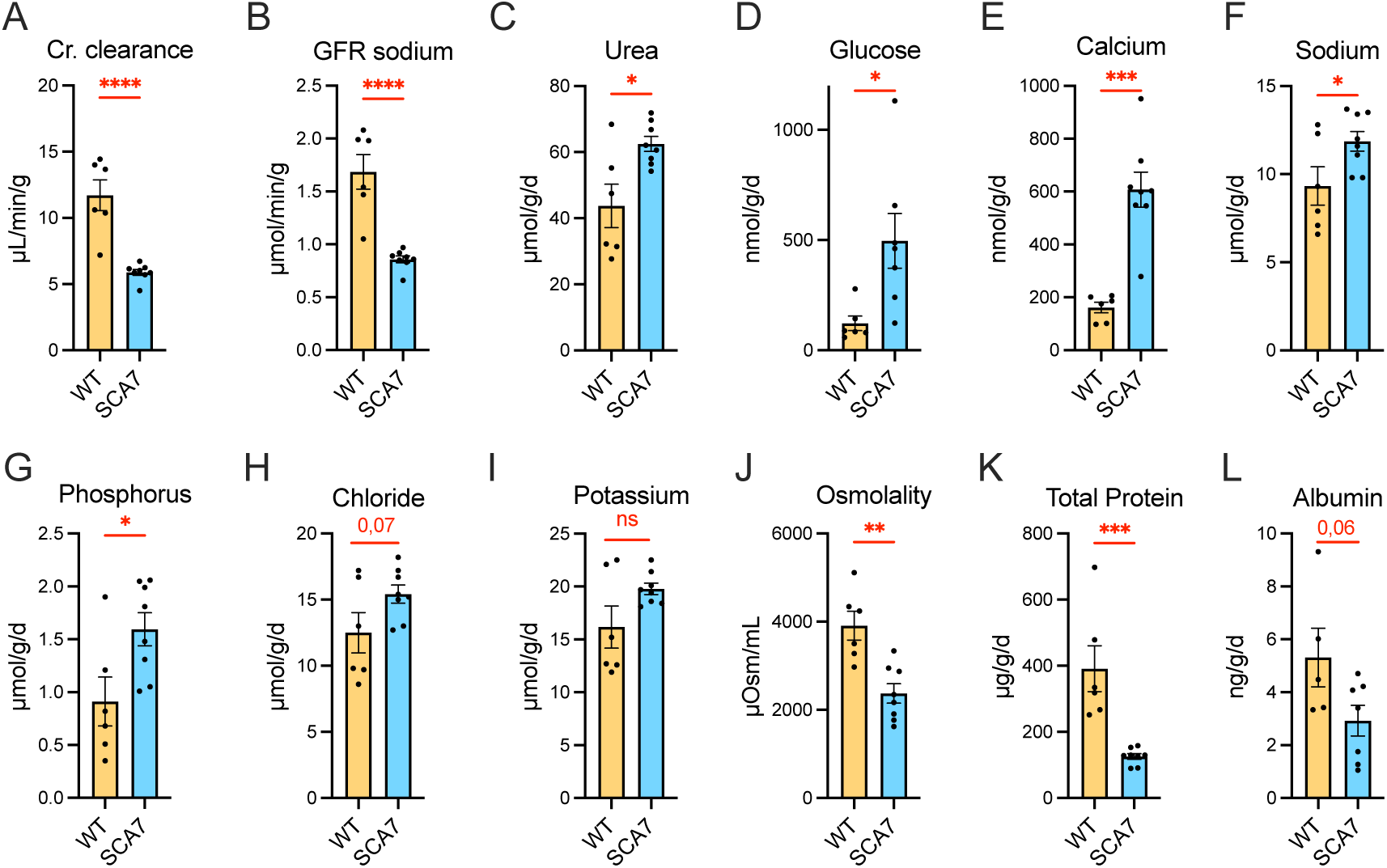
Urinary biochemistry parameters of SCA7 mice in metabolic cage. **A-B**) Creatinine (Cr.) clearance and sodium glomerular filtration rate (GFR sodium) in 24-hour urine, normalized for BW of 18-wk-old SCA7 and WT male littermates. **C-L**) Levels of 24-hour urine components normalized for BW, and calculated osmolality. Data are means ± SEM. *p* value compared WT (n= 6) versus SCA7 (n= 8) urine parameters using unpaired Student t-test. * *p* < 0.05, ** *p*< 0.01, *** *p*< 0.001, **** *p*< 0.0001. ns, not significant.

These results demonstrate that 18-wk-old SCA7 mice exhibit renal failure without proteinuria and characterized by impaired clearance of plasma waste products and defective urinary concentration. Collectively, the weight loss, reduced creatinine clearance, and the decreases in urinary osmolality and sodium concentration suggest a possible prerenal kidney injury related to sodium and water loss and extracellular volume contraction. SCA7 mice also display significant alterations in proteins, glucose and electrolyte homeostasis in fluids.

### Tubular and glomerular alterations in SCA7 mice at advanced disease stages

To examine the renal parenchyma, we conducted histological and electron microscopy analyses of kidneys from SCA7 mice at advanced disease stages. Hematoxylin-eosin staining revealed significant local hyperplasia of distal and collecting tubules in SCA7 kidneys, with an apparent increase in cellularity, as well as hypertrophy of proximal tubule cells, compared to age-matched WT controls (Figure 4A-B). Additionally, epithelial cells of the distal tubule near the renal corpuscles exhibited pyknotic nuclei, a sign of progressive cell death (Figure 4C-D). SCA7 renal corpuscles showed frequent dilation of Bowman’s capsules and the glomeruli, compared to WT (Figure 4C-D), clearly seen on electron micrographs (Figure 4E-F).

**Figure 4:**
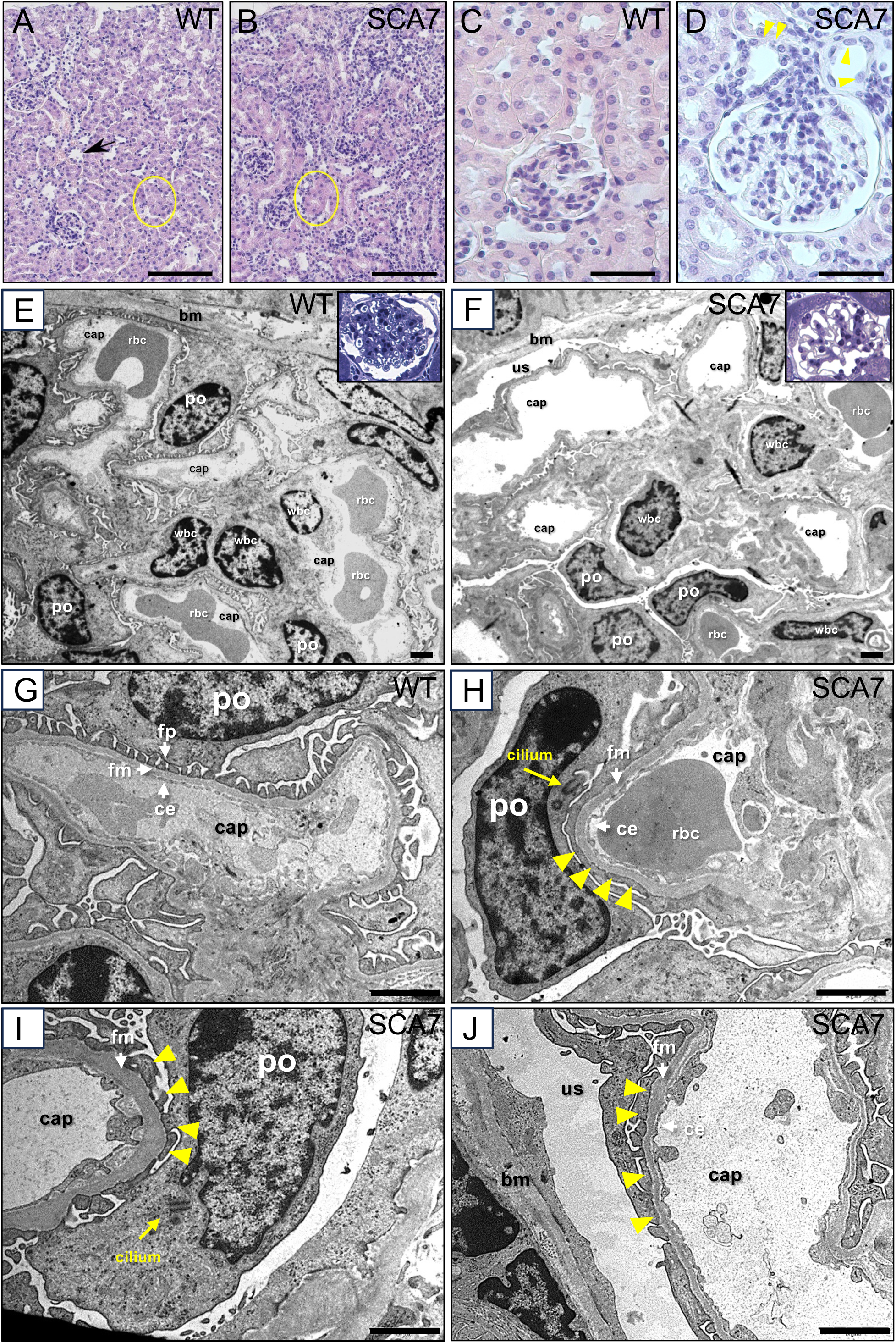
Glomerular and tubular alterations of SCA7 mice kidney. **A, B**) Hematoxylin-eosin staining of kidney sections showing hypertrophic cells in proximal tubule section (circled) of SCA7 kidney, compared to WT, and the preeminent presence of distal tubules (hyperplasia) in SCA7 tissue. Arrow in WT points to normal distal tubule section. **C, D**) Compared to WT, SCA7 Bowman’s capsule and glomerulus are dilated, and the adjacent distal tubule cells presented pyknotic nuclei (arrowheads). **E)** Glomerulus of WT mice. Basement membrane (bm), capillary loop (cap), podocytes (po), red blood cell (rbc), white blood cell (wbc). **F)** Glomerulus of SCA7 mice with dilation of capillary loop and urinary space (us). **G)** Illustration of WT podocyte foot processes (fp) that form small, regular and dense pedicels adjacent to the filtration membrane (fm) and fenestrated capillary endothelium (ce). **H, I**) SCA7 podocyte showing the abnormal presence of a cilium and effacement of pedicels, which are replaced by extended foot processes along the filtration membrane (yellow arrow heads). **J**) Illustration of pedicel effacement, replaced by extended foot processes (yellow arrow heads) in SCA7 glomerulus. Scale bar: 100 µm (A-B), 25 mm (C-D), 2 µm (E-J).

Subcellular analysis revealed that, in contrast to the regular, thin and dense foot process of WT podocytes, which forms pedicels tightly intertwined with filtration slit diaphragms at the filtration membrane (Figure 4G), SCA7 podocytes in dilated Bowman’s capsules displayed irregular and elongated foot processes, leading to significant loss of filtration slit diaphragms, a pathological feature known as pedicel effacement (Figure 4H-J). Interestingly, the cell body of SCA7 podocytes often contained a cilium (Figure 4H-I), a subcellular appendage that is normally present only in undifferentiated podocytes. In many cases, the filtration membrane in SCA7 glomeruli appeared thicker than that in WT (Figure 4I).

Together, these morphological alterations of renal tubules and glomeruli indicate that beyond a prerenal mechanism, chronic lesions may also be responsible for intrinsic renal failure.

### Hepatic dysfunction associated with altered lipid and lipoprotein metabolism

To investigate liver function, we first measured plasma levels of enzymes and other liver-released components, which can increase when liver cells are damaged. As shown in Figure 5A-C, plasma levels of alanine amino transferase (ALAT), lactate dehydrogenase (LDH) and alkaline phosphatase (ALP) were similar between SCA7 and WT mice at 30 and 43 wks. However, plasma total cholesterol (T-Chol) and triglycerides (TG) levels were significantly elevated in SCA7 mice at 30 wks (TC: +55% (p< 0.0001) and TG: +50% (p= 0.0002)) and 43 wks (TC: +50% (p< 0.0001), and TG: +100% (p< 0.0001)) (Figure 5D-E). Alterations of plasma T-Chol levels were already detected at 16 wks, particularly with a 110% increase in LDL-associated cholesterol (LDL-Chol) (p< 0.0001), while HDL-associated cholesterol (HDL-Chol) remained unchanged (Figure 5F-H). Additionally, plasma total bilirubin levels were higher in SCA7 mice at 30 wks (+60%, p= 0.0003) and 43 wks (+17%, p= 0.0193) compared to WT mice (Figure 5I).

**Figure 5:**
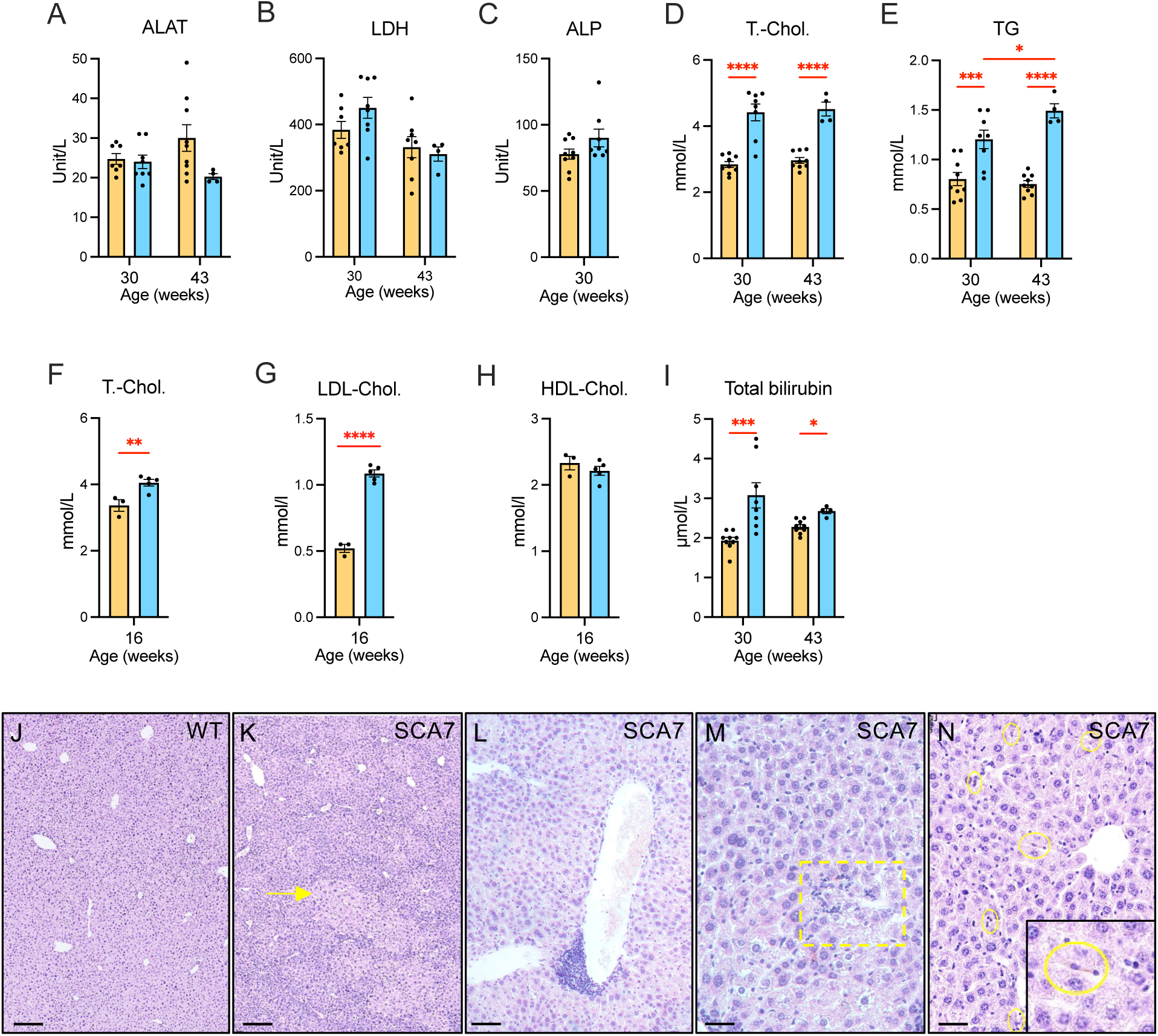
Hepatic dysfunction and lesions in SCA7 mice. **A-C**) Plasma concentration of hepatic enzymes of SCA7 (blue bar) and WT (orange bar) male littermates at 30 and 43 wks: alanine amino transferase (ALAT), lactate dehydrogenase (LDH) and alkaline phosphatase (ALP). **D-E**) Plasma concentration of total cholesterol (T-Chol) and triglycerides (TG) at 30 and 43 wks, respectively. **F-H**) Plasma concentration of T. Chol., low-density lipoprotein-associated cholesterol (LDL-Chol) and high-density lipoproteins-associated cholesterol (HDL-Chol) at 16 wks. **I**) Plasma concentration of total bilirubin at 30 and 43 wks. **J-N**) Hematoxylin-eosin staining of WT liver section (J) and SCA7 liver section showing foci of hypertrophied hepatocytes (K, yellow arrow), inflammation (L), hepatocyte cell death (M, yellow square) and pigmented Kupffer cells (N, yellow circle). Data in A-I are means ± SEM and were analyzed using unpaired Student t-test (C, F, G, H) (n= 3 WT and 5 SCA7), or the mixed-effects model (A, B, D, E, I) (30 wks: n= 9 WT and 10 SCA7; 43 wks: n= 9 WT and 4 SCA7); ALAT: F(1,24)= 3.99, p= 0.0571; LDH: F(1,23)= 0.51, p= 0.4799; T. Chol.: F(1,15)= 37.5, p< 0.0001; TG: (F(1,15)= 48.2, p< 0.0001); Total bilirubin: (F(1,15)= 13.4, p= 0.0023). * *p* < 0.05, *** *p*< 0.001, **** *p*< 0.0001. Scale bars are 200 µm (J, K), 100 µm (L), 50 µm (N, M).

Histological analysis of SCA7 liver tissues showed foci of hypertrophied hepatocytes (Figure 5J-K), inflammation (Figure 5L), hepatocyte cell death (Figure 5M) and scattered pigmented Kupffer macrophages (Figure 5N), likely reflecting red blood cell phagocytosis and heme accumulation. Therefore, SCA7 mice show early hepatic dysfunction leading to hypercholesteremia, hypertriglyceridemia, alteration in the metabolism of lipoprotein particles, together with histopathological lesions.

### Iron deficiency anemia in SCA7 mice

The elevated bilirubin levels and presumed heme accumulation in Kupffer cells suggested an increased red blood cells destruction. To investigate this, we performed hematological analysis of blood collected at 18 wks during the metabolic cage experiment. SCA7 mice showed a significant 18% reduction of red blood cells (RBC) (p= 0.008) (Figure 6A). The reduction was accompanied by hypochromia, as evidenced by significant reductions in hemoglobin concentration (HGB) (-21%, p= 0.008), hematocrit percentage (HCT) (-18%, p= 0.011), and mean corpuscular hemoglobin concentration (MCHC) (-24%, p= 0.029), while the mean corpuscular volumes (MCV) of RBC were normal (Figure 6B-E). Similar findings were also observed at 30 and 43 wks (Supplementary figure S2A-D). White blood cells (WBC) and platelets (Plt) counts were comparable between SCA7 and WT mice at 18 wks (Figure 6F-G). However, Plt counts were significantly elevated in SCA7 mice at 30 and 43 wks (Supplementary figure S2G), indicating late thrombocytosis.

**Figure 6:**
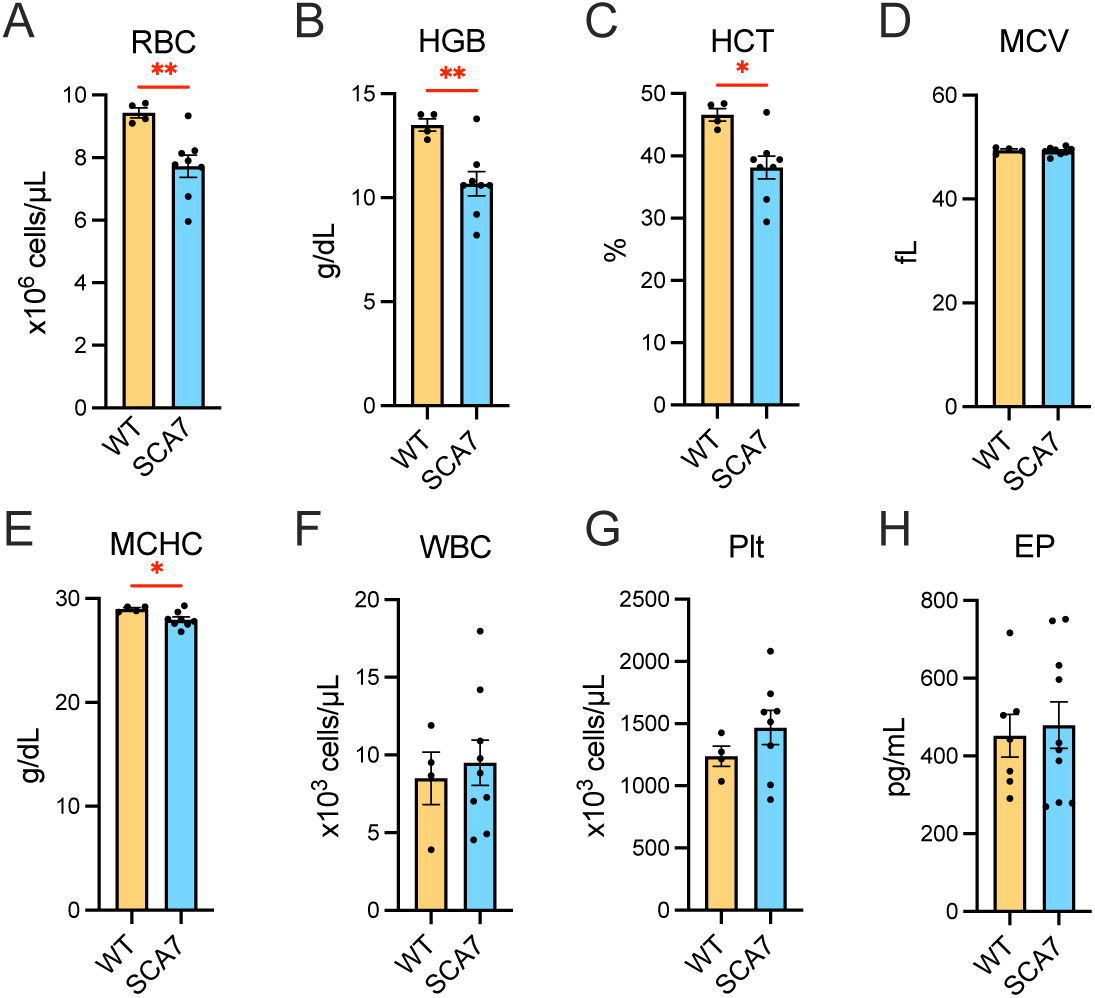
Early alteration of red blood cells. **A)** Red blood cells (RBC) count in SCA7 and WT male littermates. **B)** Hemoglobin (HGB) concentration. **C)** Percentage of hematocrit (HCT) in plasma. **D)** The mean corpuscular volume (MCV) of red blood cells. **E)** The mean corpuscular hemoglobin concentration (MCHC). **F)** White blood cells (WBC) count. **G)** Platelet (Plt) count. **H)** Plasma concentration of erythropoietin (EP). Mice were analyzed at 18 wks. Data are means ± SEM and were analyzed using the unpaired Student t-test (18 wks: n= 4 WT and 8 SCA7). * *p* < 0.05, ** *p* < 0.01.

Given that the reduced RBC counts coincided with renal dysfunction, we measured the plasma erythropoietin level, a hormone secreted by the kidney to stimulate erythropoiesis. The erythropoietin level was comparable in WT and SCA7 mice at 18 wks (Figure 6H), suggesting inability of kidney to compensate the reduced RBC. We then explored the possibility that the reduction in RBC count, size and hemoglobin content could come from impaired iron metabolism. Plasma iron levels were significantly decreased in SCA7 mice with a 25% reduction at 16 wks (p= 0.0027) and 17% reduction at 43 wks (p= 0.0022) (Figure 7A). Consistently, the transferrin-iron saturation coefficient (TSC) decreased by 34% at 16wks (p= 0.0005) and 23% at 43 wks (p= 0.0006), even if there was a slight increase in plasma transferrin (TF) levels (+16%, p< 0.0001) at this age (Figure 7B-C). Therefore, iron deficiency anemia represents an early manifestation in SCA7 mice.

**Figure 7.**
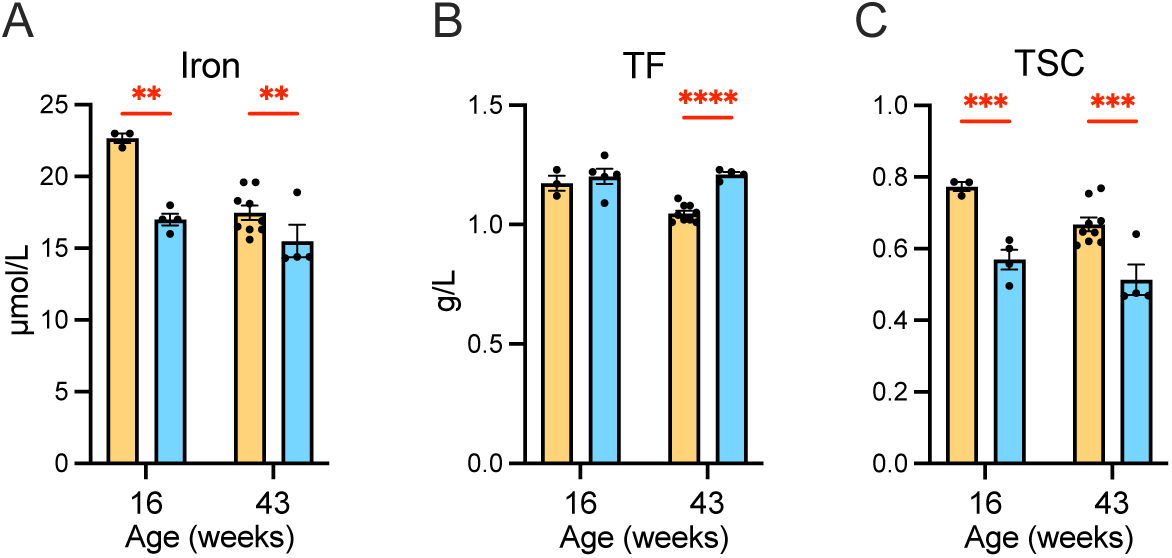
Early alteration of iron metabolism. **A)** Plasma concentration of iron. **B)** Plasma concentration of transferrin (TF). **C)** Calculated transferrin saturation coefficient (TSC). Two different mice cohorts were analyzed at 16 and 43 wks. Data are means ± SEM and were analyzed using the unpaired Student t-test (16 wks: n= 3 WT and 4 SCA7; 43 wks: 9 WT and 4 SCA7). ** *p* < 0.01, *** *p*< 0.001, **** *p*< 0.0001.

### Early cardiac dysfunction in SCA7 mice

To determine whether polyuria-induced hypovolemia could affect cardiac function in SCA7^140Q/5Q^ mice, we performed echocardiographic analysis on males at 11 and 26 wks of age, corresponding to onset and moderate stages of the pathology [25]. At these ages, SCA7 males exhibited a 10% (p= 0.040) and 25% (p< 0.0001) reduction in body weight, respectively, compared to age-matched WT littermates (Figure 8A). When normalized to body weight, the cardiac output in 11-wks-old SCA7 mice was 31% lower than in WT mice (p= 0.041) (Figure 8B), primarily due to a decreasing trend in stroke volume (-23%, p= 0.07) (Figure 8C). At 26 wks, most SCA7 mice continued to show lower cardiac output and stroke volume, though the overall differences were not statistically significant due to high interindividual variability. Heart rate remained comparable between groups at both ages (Figure 8D). Overall, the data suggest that hypovolemia contributes to reduced cardiac output in SCA7 mice without triggering compensatory tachycardia.

**Figure 8.**
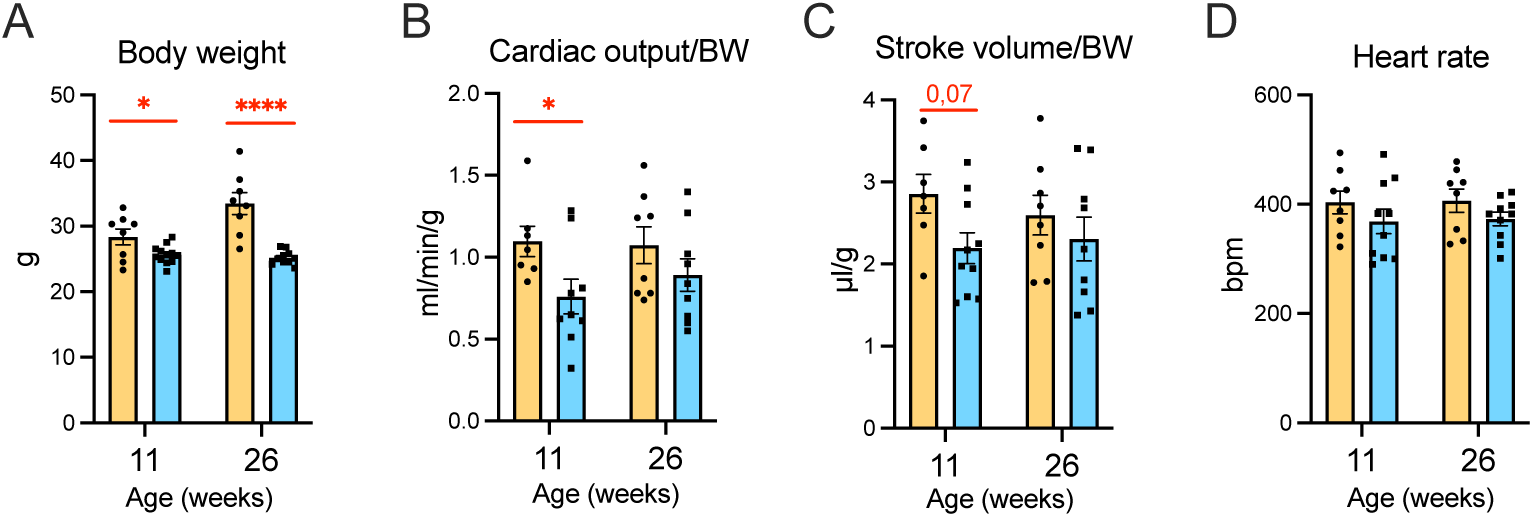
Echocardiography reveals a reduced cardiac output in SCA7 mice. **A)** Body weight (BW) of SCA7 (blue bar) and WT (orange bar) male littermates at 11 and 26 wks. **B)** Cardiac output normalized for BW. **C)** Stroke volume normalized for BW. **D)** Heart rate (bpm, beats per minute). Data are means ± SEM and were analyzed using the mixed-effects model (11 wks: n= 8 WT and 12 SCA7; 26 wks: n= 8 WT and 10 SCA7). (BW: F (1,18) = 20.6, p= 0.0003; Cardiac output/BW: F (1,17) = 5.2, p= 0.036; Stroke volume/BW: F (1,17) = 3.3, p= 0.085; Heart rate: F (1,18) = 2.7, p= 0.116). * *p* < 0.05, **** *p*< 0.0001.

## DISCUSSION

### Peripheral pathology in SCA7

Most research on SCA7 has focused on the primary neurological symptoms, particularly cerebellar and retina degeneration. However, ATXN7 is ubiquitously expressed, and polyQ-expanded ATXN7 aggregates have been detected in peripheral tissues, such as kidney and heart, especially in patients with large expansions [10,32]. Our findings extend these observations by showing that adult SCA7^140Q/5Q^ mice develop early renal, hepatic, cardiac dysfunctions that give rise to broad metabolic disturbances - including uremia, altered calcium-phosphorus balance, dyslipidemia, anemia and inflammation – that emerge prior detectable motor deficits. These systemic changes likely contribute to comorbidity, growth deficits, general weakness and premature mortality, underscoring the need to consider extra-neural pathology when designing therapeutic strategies.

### Renal pathology and tubular dysfunction

In human infantile SCA7, renal biopsies have revealed diverse pathological alterations, including focal segmental glomerulosclerosis, cytoplasmic inclusion in podocytes, protein resorption droplets in tubule cells and other unspecified glomerular and tubular changes [11,13]. Clinically, these early-onset cases were marked by elevated plasma urea and creatinine levels together with proteinuria. Similarly, we found that 18-wk-old SCA7 mice exhibited elevated plasma urea and creatinine, but their renal phenotype was distinct: instead of proteinuria, they displayed excessive production of dilute urine containing abnormally high levels of calcium and glucose. This suggests primary disturbances of both proximal and distal tubular function. The polyuria-polydipsia syndrome, in absence of proteinuria, may reflect impaired vasopressin (antidiuretic hormone) signaling, as both distal tubular cells in the collecting duct, which express the V2 vasopressin receptor, and hypothalamic vasopressin neurons express ATXN7 and contain mutant ATXN7 aggregates [10,33]. Our histology indeed revealed epithelial cell loss in distal tubules as well as hypertrophy of proximal tubule cells and dysplasia of distal and collecting tubules, pointing to a progressive tubular pathology. Although glomerular functions appeared preserved at early disease stage, later changes included pedicel effacement, a hallmark of podocyte injury, raising the possibility of delayed-onset proteinuria.

### Loss of podocyte differentiation

A striking finding was the presence of primary cilia in podocytes of adult SCA7 mice. Since cilia are normally lost during terminal podocyte differentiation [34], their presence suggests impaired maturation. Together with pedicel effacement, this phenotype parallel defects in SCA7 photoreceptors, where polyQ-expanded ATXN7 disrupts transcriptional programs essential for maintaining outer segments, a highly differentiated structure essential for phototransduction [23,26]. We propose that a similar mechanism—failure to maintain cell identity genes—underlies podocyte dedifferentiation in SCA7. This mechanism could be especially relevant to congenital/infantile forms, where very large expansions may impair podocyte maturation during development, leading to early renal dysfunction.

### Hepatic and cardiac contributions

Hepatic alterations included increased production of lipoproteins, cholesterols, and triglycerides, likely reflecting chronic inflammation and altered plasma homeostasis. These changes were accompanied by elevated bilirubin and reduced iron, which may contribute to anemia. Cardiac dysfunction emerged even earlier: reduced cardiac output and stroke volume was observed by 11 wks, preceding neurological decline. This impairment may worsen renal function via reduced perfusion, consistent with a cardiorenal syndrome. In patients, severe forms of SCA7 are often associated with hypertrophic cardiomyopathy and congenital heart defects [6,11]. While our echocardiography setup was not capable of detecting structural anomalies in SCA7 mice, prior studies in polyQ models show that aggregate-prone proteins are intrinsically toxic to cardiomyocytes [35,36]. Thus, cardiac pathology in SCA7 may reflect both intrinsic toxicity of mutant ATXN7 and secondary systemic effect.

### Systemic and therapeutic implications

The metabolic profile of SCA7 mice—anemia, dyslipidemia, altered calcium–phosphate balance— likely arises from the interplay of renal, hepatic, cardiac, and possibly CNS dysfunction. Disentangling primary from secondary events will require additional longitudinal studies. Nonetheless, our findings align with growing evidence that peripheral pathology is a critical component of polyQ diseases [30,31]. Importantly, peripheral organs are accessible therapeutic targets: interventions that ameliorate muscle or cardiac dysfunction improved outcomes in SCA1 and HD mouse models [37,38]. Similarly, peripheral biomarkers, such as early metabolic changes in SCA7 mice, could aid in disease monitoring and treatment evaluation.

## Conclusion

Our study demonstrates that peripheral organ dysfunction is an early and integral component of SCA7 pathology in mice, preceding classical neurological symptoms. The identification of renal tubular defects, podocyte dedifferentiation, hepatic dysregulation, and cardiac failure highlights the systemic nature of SCA7. These findings broaden the view of SCA7 from a neurodegenerative disease to a multi-organ disorder, with important implications for biomarker discovery and therapeutic strategies.

## MATERIALS AND METHODS

### Mouse information

SCA7^140Q/5Q^ knock-in mice were maintained on a C57Bl/6J background and bred at the Mouse Clinical Institute under controlled environmental conditions: temperature of 21 ± 2 °C, 60 ± 5% relative humidity, and 12-hour light-dark cycle. Food and water were provided *ad libitum*. Genotyping was performed by PCR as described previously [28]. The expansion size ranged from 140 to 150 CAG repeats in exon1 of the *Atxn7* locus. Unless otherwise specified, analyses were performed on male WT and SCA7^140Q/5Q^ littermates.

### Metabolic cage studies, sample collection and analysis

Eighteen-week-old WT and SCA7 males were individually housed in metabolic cages (Tecniplast, Italy), designed for separate urine and feces collection. Mice maintained under controlled temperature and humidity, with a 12-hour light-dark cycle and free access to water and a standard chow (D04, SAFE Villemoisson-sur-Orge, France). Animal were acclimatized to the cages for 3 days before urine collection to minimize stress. Urines were collected over a 24-hour period, measured for total volume and stored at -80°C. Water consumption was recorded. At the end of the study, mice were weighed and blood was collected by the temporo-mandibular veinpuncture. Plasma was obtained by centrifugation at 5,000 rpm for 15 min at 4°C. Plasma and urine analysis were performed on an OLYMPUS AU-480 automated laboratory workstation (Beckmann Coulter, US) using manufacturer-supplied kits and controls. The following parameters were quantified: urea, creatinine, total protein, albumin, glucose, calcium, sodium, chloride, potassium, phosphorus, iron, ferritin, transferrin, erythropoietin. Urinary osmolality was calculated using the formula: ((µmol/mL Na + µmol/mL K) x 2) + µmol/mL urea. Additional plasma biochemistry analyses were performed on 16-, 30- and 43-wk-old animals, and included total cholesterol, LDL- and HDL-cholesterol, triglycerides, LDH, ALAT, ALP and total bilirubin. Complete blood count, HGB and MCV, HCT and MCHC were assessed using an Advia 120 Vet (Siemens Healthcare Diagnostic, Saint-Denis, France).

### Echography Image acquisition

Mice were anaesthetized using inhaled isoflurane (3% for induction, 1.5-2% for maintenance) and positioned supine on a heated imaging platform (37°C). Eye lubricant was applied, and a rectal probe was used to monitor body temperature and heart rate. The heart rates were maintained above 450 bpm by adjusting anesthesia as needed. Echography was performed with Vevo-2100 system (FUJIFILM VisualSonics, Inc., Toronto, Canada) using a 30 MHz linear array transducer and MS400 for heart imaging.

For heart imaging, standard parasternal long-axis and short axis views at the mid-papillary level were obtained to assess the cardiac morphology, ventricular function, and hemodynamics. Stroke volume and cardiac output were calculated using Doppler-derived velocity–time integral (VTI) measurements and left ventricular outflow tract (LVOT) diameter. Following imaging, mice were allowed to recover with supplemental oxygen and returned to their cages

### Histology and electron microscopy analysis

Mice were weighed and sacrificed for necropsy. Liver, kidneys, heart, and spleen were weighed, and fixed in paraformaldehyde, embedded in paraffin, sectioned (5 µm), and stained with hematoxylin-eosin.

For electron microscopy, animals were perfused with 5 mL PBS followed by 10 mL 4% paraformaldehyde. Tissues were fixed in 2.5% paraformaldehyde /2.5% glutaraldehyde in 0.1M cacodylate buffer (pH 7.4), post-fixed in 1 % osmium tetroxide for 1-hour at 4 °C, and dehydrated in graded alcohol (50-100 %) and propylene oxide for 30 min each, embedded in Epon 812. Semi-thin (2 µm) sagittal sections were cut with Leica Ultracut UCT ultramicrotome, stained with toluidine blue, and examined by light microscopy. Ultra-thin (70 nm) sections were cut and contrasted with uranyl acetate and lead citrate and examined at 70 kV with a Morgagni 268D electron microscope (FEI Electron Optics, Eindhoven, the Netherlands). Images were captured digitally by Mega View III camera (Soft Imaging System).

### Statistical analysis

Data were analyzed using GraphPad Prism 8. Normality was assessed by the Shapiro-Wilk’s test and outlier by the Grubbs test (significance level at 0,05). Differences between groups were evaluated with unpaired Student’s *t*-test or mixed-effect models, as appropriate. Significance was established at p< 0,05. Results are expressed as mean ± SEM unless otherwise indicated. Further information is indicated in the figure legends. Datasets used and/or analyzed during the current study are available from the corresponding author upon reasonable request.

## Author contributions

Conceptualization, Resources, project management: Y.T. Investigation: GAB, MFC, NM, CW, ANC. Formal analysis: LM, GAB, MFC, NM, MS, BM. Data curation: GAB, YT. Visualization: NM, YT. Writing original draft: YT. Text editing: LM, BM. All authors have read and approved the final manuscript.

## Institutional Review Board Statement

All animal experiments complied with French national laws on laboratory animal welfare and the guidelines of the Federation of European Laboratory Animal Science Associations, based on European Union Legislation (Directive 2010/63/EU). Facilities were registered for animal housing and experimentation, and investigators held certificates authorizing experimentation issued by the governmental veterinary office. Protocols were approved by local Animal Ethics Committees under the supervision of the French Ministry of Higher Education, Research and Innovation (agreement numbers 5149-20160422171329).

## Informed Consent Statement

Not applicable.

## Data Availability Statement

Data that support the findings of this study are available from the corresponding authors on request.

## Acknowledgments

We thank the members of the Mouse Clinical Institute of Illkirch for mouse phenotyping support; Josiane Hergueux and Jean-Luc Weickert for technical assistance at the electron microscopy platform; Nathalie Daigle for critical reading of the manuscript and fruitful discussion.

---

**Supplementary figure S1: Plasma biochemistry of SCA7 mice at 16, 30 and 43 wks of age.**

**A**) Body weight of wild type (WT, orange) and SCA7 (blue bar) males at 30 and 43 wks.

**B-K**) Concentration of plasma components at 30 and 43 wks.

**L-N**) Concentration of plasma components WT and SCA7 males and females at 16 wks of age. Data are means ± SEM and were analyzed using unpaired Student t-test at 16 wks (n= 3 WT and 5 SCA7 males, n= 4 WT and 2 SCA7 females), and the mixed-effects model at 30wks (n= 9 WT and 8 SCA7 mice) and 43 wks (n= 9 WT and 4 SCA7 mice) (BW: F(1,15)= 57.14, p< 0.0001; Urea: F(1,15)= 2.41, p= 0.141; Creatinine: F(1,15)= 97.4, p< 0.0001; Total proteins: F(1,26)= 13.4, p= 0.011; Albumin: F(1,15)= 41.9, p< 0.0001; Glucose: F(1,15)= 13.25, p= 0.0024; Calcium: F(1,15)= 57.7, p< 0.0001; Sodium: F(1,15)= 61.6, p< 0.0001; Chloride: F(1,26)= 101, p< 0.0001); Phosphorus: F(1,15)= 0.9849, p= 0.3367; Potassium: F(1,15)= 0.0283, p= 0.8687) * *p* < 0.05, ** *p*< 0.01, *** *p*< 0.001, **** *p*< 0.0001.

**Supplementary figure S2: Alteration of red blood cells and iron metabolism at late disease stages.**

**A)** Red blood cells (RBC) count in SCA7 (blue bar) and WT (orange bar) male littermates at 30 and 43 wks.

**B)** Hemoglobin (HGB)concentration.

**C)** Percentage of hematocrit (HCT).

**D)** Mean corpuscular hemoglobin concentration (MCHC).

**E)** Mean corpuscular volume (MCV) of red blood cells.

**F)** White blood cells (WBC) count.

**G)** Platelet (Plt) count.

Data are means ± SEM and were analyzed using the mixed-effects model (30wks: n= 9 WT and 8 SCA7 mice; 43 wks: n= 9 WT and 4 SCA7 mice). (RBC: F (1, 26) = 48.3, p< 0.0001; HGB: F (1, 26) = 140, p< 0.0001; HCT: F (1, 26) = 88.5, p< 0.0001; MCHC: F (1, 26) = 81.3, p< 0.0001; MCV: F (1, 15) = 5.1, p= 0.0393; WBC: F(1,15)= 3.56, p=0.0785; Plt: F (1, 15) = 62.24, p< 0.0001). ** *p* < 0.01, *** *p*< 0.001, **** *p*< 0.0001.

