## Supplemental Figures S1 and S2 for "Cardiorenal and hepatic dysfunctions underlie metabolic alterations in the spinocerebellar ataxia type 7 mice"

### Supplementary Figure S1

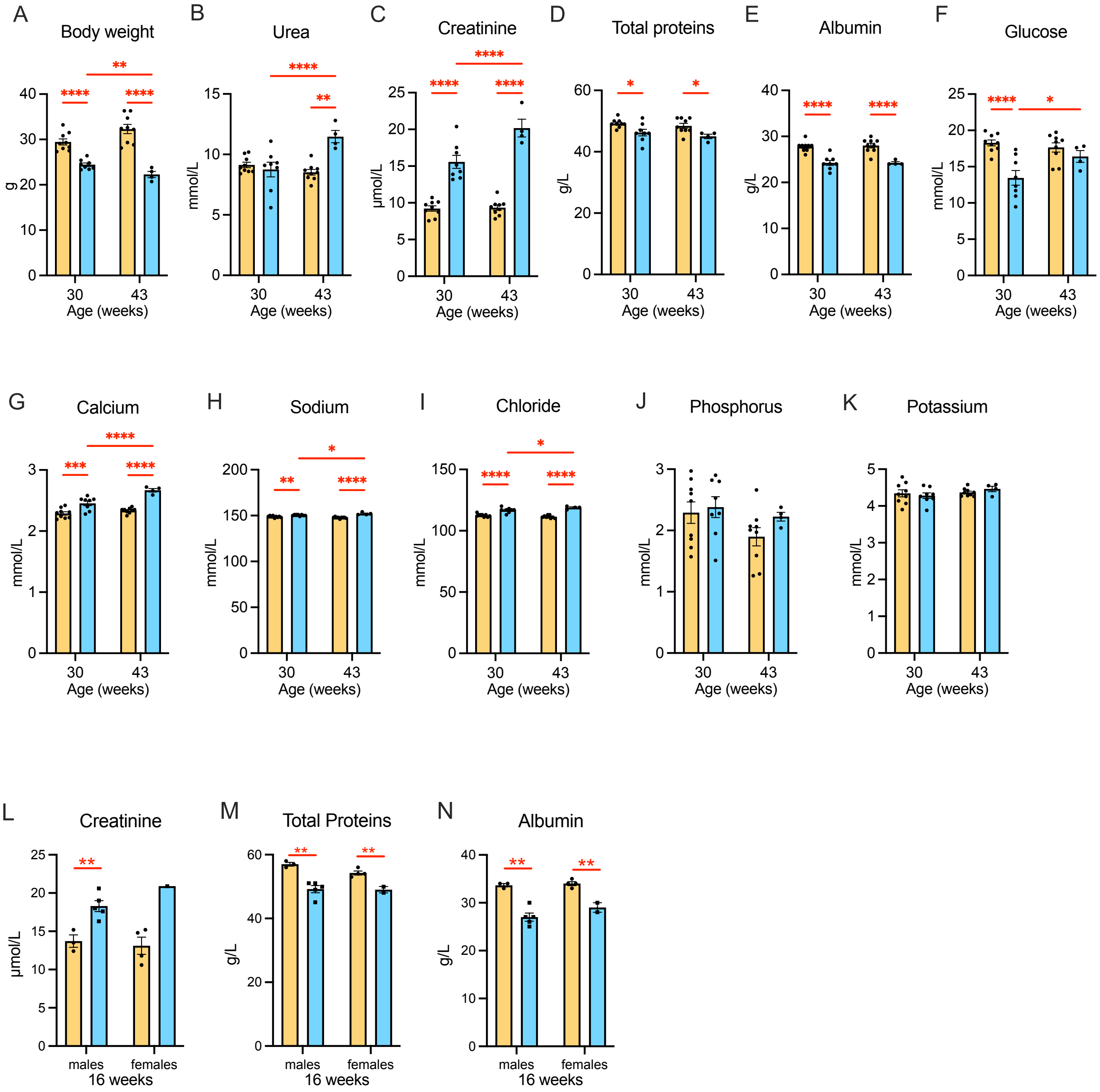


**Figure S1**: Plasma biochemistry of SCA7 mice at 16, 30 and 43 wks of age. **(A)** Body weight of wild type (WT, orange) and SCA7 (blue bar) males at 30 and 43 wks. **(B-K)** Concentration of plasma components at 30 and 43 wks. **(L-N)** Concentration of plasma components WT and SCA7 males and females at 16 wks of age. Data are means ± SEM and were analyzed using unpaired Student t- test at 16 wks (n= 3 WT and 5 SCA7 males, n= 4 WT and 2 SCA7 females), and the mixed-effects model at 30wks (n= 9 WT and 8 SCA7 mice) and 43 wks (n= 9 WT and 4 SCA7 mice) (BW: $F(1,15)= 57.14$, $p< 0.0001$; Urea: $F(1,15)= 2.41$, $p= 0.141$; Creatinine: $F(1,15)= 97.4$, $p< 0.0001$; Total proteins: $F(1,26)= 13.4$, $p= 0.011$; Albumin: $F(1,15)= 41.9$, $p< 0.0001$; Glucose: $F(1,15)= 13.25$, $p= 0.0024$; Calcium: $F(1,15)= 57.7$, $p< 0.0001$; Sodium: $F(1,15)= 61.6$, $p< 0.0001$; Chloride: $F(1,26)= 101$, $p< 0.0001$); Phosphorus: $F(1,15)= 0.9849$, $p= 0.3367$; Potassium: $F(1,15)= 0.0283$, $p= 0.8687$) * $p < 0.05$, ** $p< 0.01$, *** $p< 0.001$, **** $p< 0.0001$.

### Supplementary figure S2

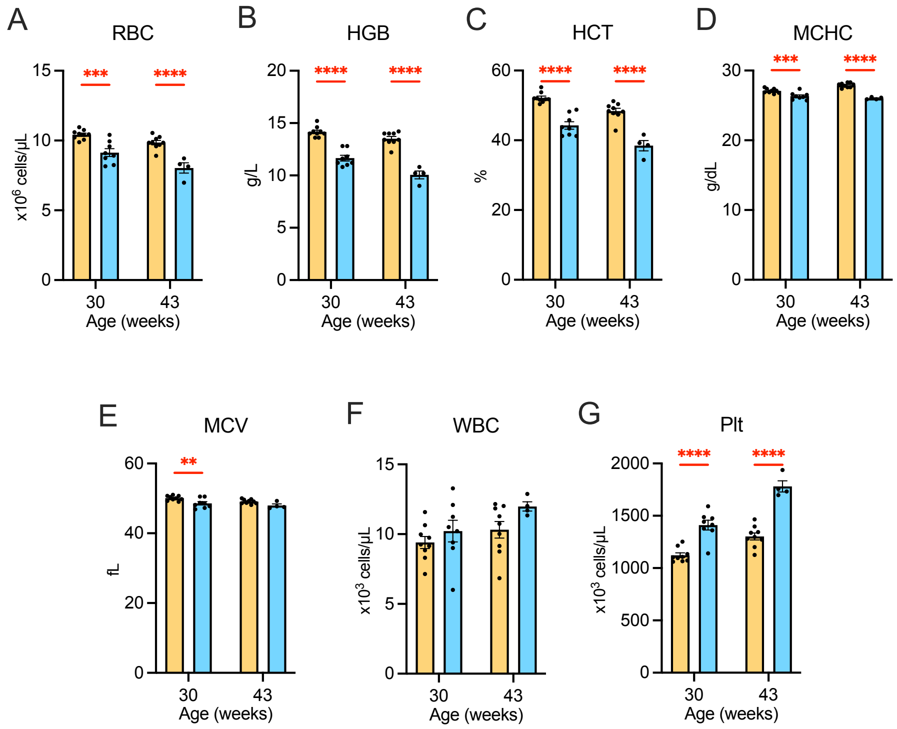


**Figure S2**: Alteration of red blood cells and iron metabolism at late disease stages. (**A**) Red blood cells (RBC) count in SCA7 (blue bar) and WT (orange bar) male littermates at 30 and 43 wks. (**B**) Hemoglobin (HGB)concentration. (**C**) Percentage of hematocrit (HCT). (**D**) Mean corpuscular hemoglobin concentration (MCHC). (**E**) Mean corpuscular volume (MCV) of red blood cells. (**F**) White blood cells (WBC) count. **G**) Platelet (Plt) count. Data are means ± SEM and were analyzed using the mixed-effects model (30wks: n= 9 WT and 8 SCA7 mice; 43 wks: n= 9 WT and 4 SCA7 mice). (RBC: F (1, 26) = 48.3, p< 0.0001; HGB: F (1, 26) = 140, p< 0.0001; HCT: F (1, 26) = 88.5, p< 0.0001; MCHC: F (1, 26) = 81.3, p< 0.0001; MCV: F (1, 15) = 5.1, p= 0.0393; WBC: F(1,15)= 3.56, p=0.0785; Plt: F (1, 15) = 62.24, p< 0.0001). ** $p < 0.01$, *** $p < 0.001$, **** $p < 0.0001$.
